# The Effect of Electrical Resonance in Neurons upon the Instability of Electrical Nerve Stimulations

**DOI:** 10.1101/2023.01.26.525813

**Authors:** Shoujun Yu, Wenji Yue, Tianruo Guo, Yonghong Liu, Yapeng Zhang, Sara Khademi, Tian Zhou, Zhen Xu, Bing Song, Tianzhun Wu, Fenglin Liu, Yanlong Tai, Xuefei Yu, Hao Wang

## Abstract

Repetitive electrical nerve stimulation can induce a long-lasting perturbation of the axon’s membrane potential, resulting in unstable stimulus-response relationships. Despite being observed in electrophysiology, the precise mechanisms underlying stimulus-induced instability is still an open question due to the lack of a proper theoretical model. This study proposes a new method based on a Circuit-Probability theory to reveal the interlinkages between the electrical resonance of neurons and the instability of neural response. Supported by in vivo investigations, this new model predicts several key characteristics of stimulus-induced instability and proposes a stimulation method to minimize the instability. This model provides a powerful tool to improve our understanding of the interaction between the external electric field and the complexity of the biophysical characteristics of axons.

## Introduction

Function electrical stimulation assists purposeful movement by increasing the plasticity for motor function. By applying electrical stimulation to the paralyzed muscles, electrical nerve stimulation makes the coordinated muscle contract in a sequence that allows paralyzed patients to perform tasks such as standing, walking, or grasping a key ^[1–7]^. However, more detailed neurological mechanisms underlying electrical stimulation are still unclear. For example, the stimulus-dependent instability of neural response is a well-known phenomenon. The axon’s excitability has a non-monotonic fluctuation with repetitive stimulation. This unstable excitability was observed from single-cell-based patch-clamping electrophysiology ^[8]^ and muscular response to FES ^[9–11]^. It is widely believed that the origin of this instability is the fluctuation of the axon’s threshold voltage. Thus, this phenomenon is also called “threshold fluctuation” or “excitability fluctuation” in previous studies ^[9–13]^.

Since this instability will affect the FES performance of the clinical treatment of diseases by electrical nerve stimulations, it concerned the researchers in this area. Many experimental studies have been conducted to investigate more detailed biological mechanisms in order to minimize the unstable effect ^[8–35]^. Major experimental observations and concluded principles of threshold fluctuation are summarized below.

1. The threshold fluctuation is caused by the perturbation of the membrane potential of the axon, where the induced membrane potential perturbation can last more than 100 ms, longer than the refractory period of the axonal action potential ^[36–42]^. Thus, it can be excluded that the neural refractory period is the major factor contributing to the stimulus-dependent instability in stimulations [9].
2. The observed membrane potential perturbation happens in the vicinity of the stimulating electrode ^[10]^. Therefore, although the EMG (Electromyography) signal is involved in evaluating the change of excitability for most relevant experiments, the muscle activation is irrelevant to the instability ^[9–11]^.
3. Subthreshold stimulations can also induce threshold fluctuation. Thus, the membrane potential perturbation happens as long as local electrical stimulations are applied, whether the action potential fires or not ^[9–11]^.
4. The relationship between the electrical stimulation parameter and the change in excitability is definitive ^[8–11, 13–14, 16, 18–21, 23–26, 32–34, 43]^. Previous studies assumed noise as the origin of threshold fluctuation ^[44, 45]^, which contradicts the experimental observation.

However, the precise mechanisms underlying the observed stimulus-dependent instability remain unclear due to the lack of a proper theoretical model. In this study, we conducted both in vivo and in silico investigations to better understand this question. Our theoretical model based on a Circuit Probability theory ^[46]^ allows us to explore how the stimulation parameters affect the instability. It reveals that the electrical resonance of neurons is the major factor to determine the instability of electrical stimulations. Meanwhile, our study provides a computational tool to better characterize neurological mechanisms underlying electrical stimulation.

## Method

### 1. Animals Preparation

Male Sprague-Dawley rats (∼300g) were used in experiments. Rats were housed and cared for in compliance with the guidelines of Institutional Animal Care and Use Committee (IACUC) and were humanely euthanized after the experiment. The rat was placed in a transparent acrylic box and anesthetized with isoflurane (Iflurin, RingPu, China; R500-Series, RWD Life Science, China). Observe the paw retraction reflex and breathing rate to estimate the depth of anesthesia. After deep anesthesia, the rat was placed on a heating pad to maintain the body temperature at 37℃ and worn an anesthesia mask during the experiment to ensure deep anesthesia. Remove the fur on the legs, disinfect the surgical area with 75% ethanol, and then expose and locate the sciatic nerve.

### 2. The instability in electrical nerve stimulations

We evaluated the instability by recording the patterns of kicking force of a rat’s leg under sciatic nerve stimulation in vivo. The experimental setup is shown in Figure 1(a). A homemade flexible neural probe (Figure 1(b)) with five channels (Figure 1(c)) connected with an FPC (Flexible Printed Circuit) connector was implanted on the sciatic nerve, shown in Figure 1(c). Since two branches of the sciatic nerve, the tibial nerve and the common peroneal nerve, control the opposite kicking, one branch was cut in the downstream location to ensure a one-directional kicking. When an electrical stimulus (STG 4008, Multi-Channel Systems GmbH, Germany) was applied to the neural probe, a kicking force (forward or backward, depending on which branch was cut off) was recorded by the force gauge (ZL-X10 & ZL-620, Anhui Yaokun Biotechnology, China) connected to the rat’s leg with a wire.

**Figure 1.**
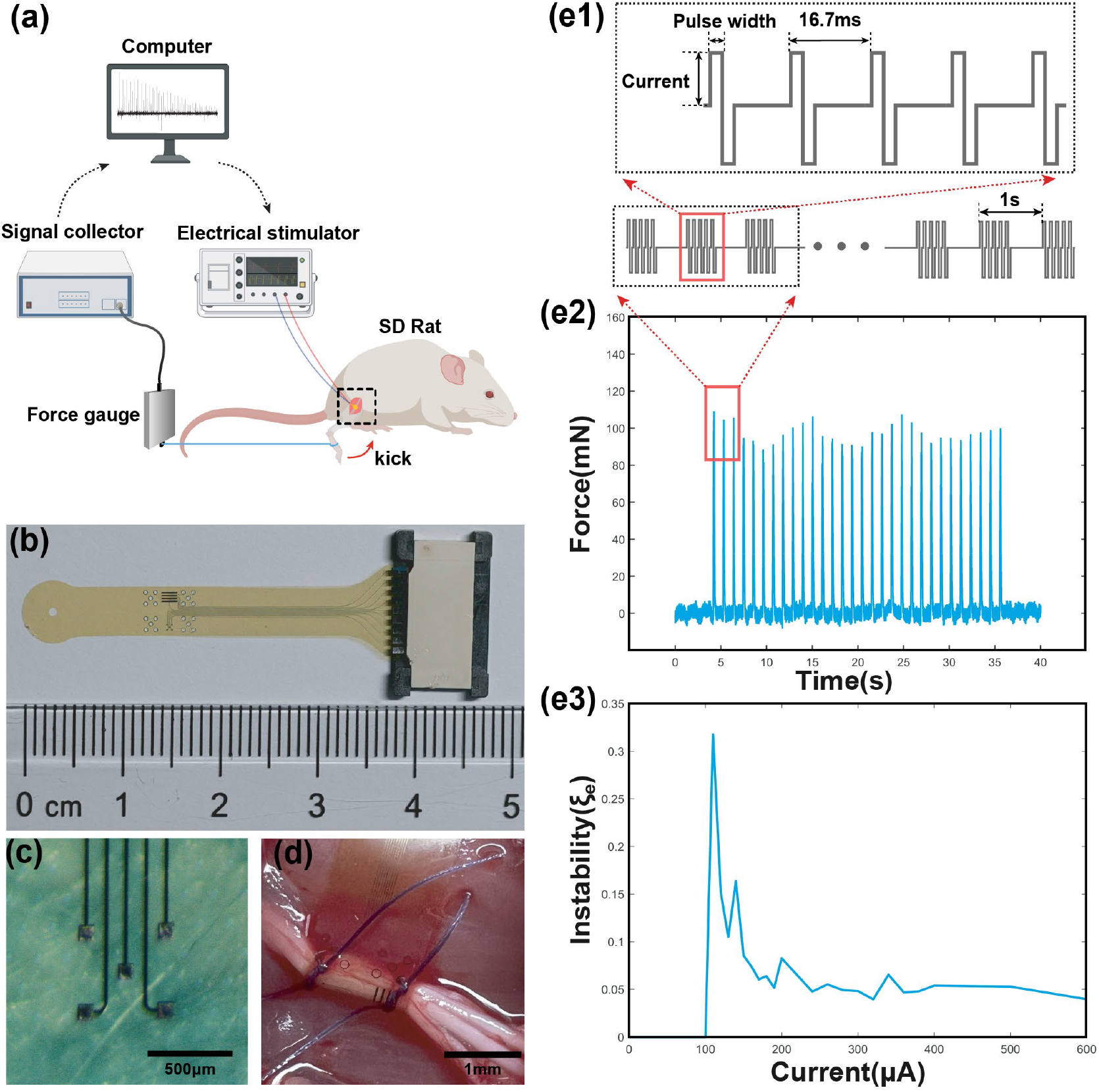
The testing setup, stimulation protocol and the result sample. (a) The testing setup for measuring the force generated by sciatic nerve stimulations; (b) The fabricated flexible neural probe used for sciatic nerve stimulations; (c) The electrode pads on the neural probe for stimulations; (d) The flexible neural probe implanted to the sciatic nerve; (e1) The stimulation protocol. Each stimulation train contains 5 current pulses with 16.7 ms as latency. The latency between each train is 1 s. (e2) The sample of measured force showing non-monotonous fluctuation; (e3) A sample of the instability curve of the measured force, defined as 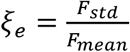 by changing the current amplitudes.

The stimulation protocol is shown in Figure 1(e). Each stimulation train contains five pulses with 16.7 ms as the latency (Figure 1(e1)). Each train generates one force pulse with a period of 1s. The detailed current waveforms, current amplitudes, and pulse widths will be provided for each test are shown in Table 2.

**Table 1.**
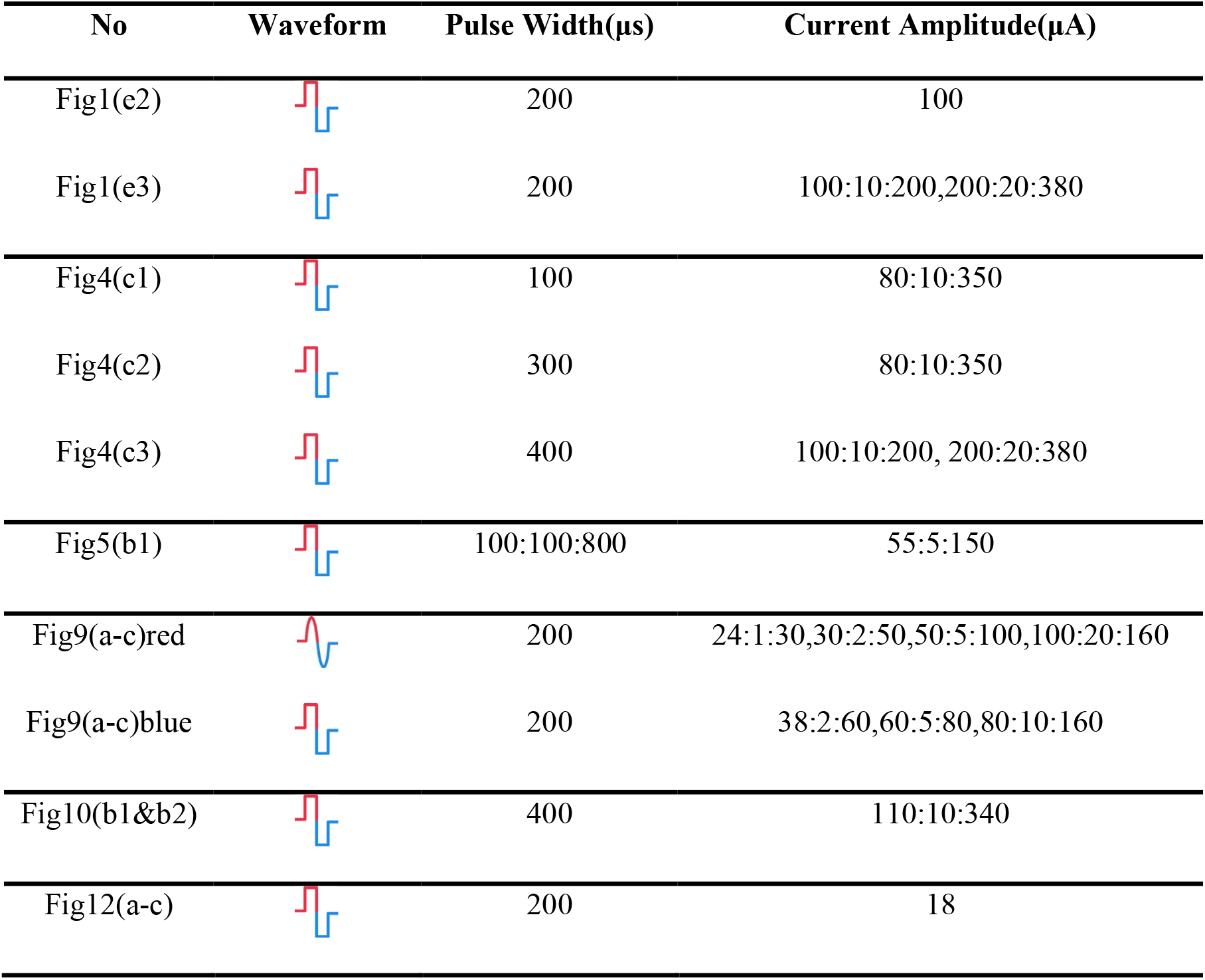
Parameters of *in-vivo* experiments.

**Table 2.**
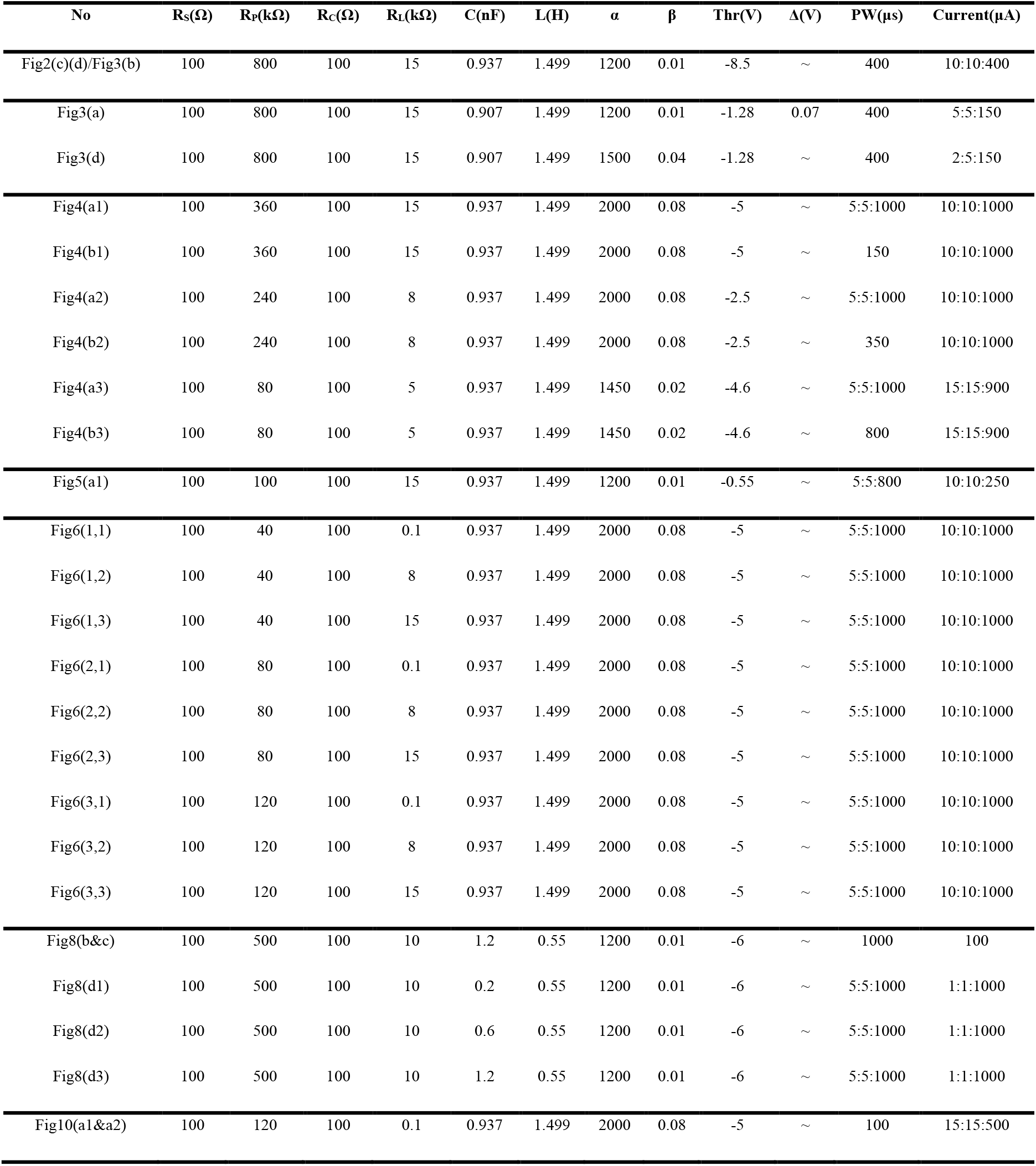
Parameters of modeling.

Figure 1(e2) shows a typical sample of the measured force with specific testing parameters (Table 2-e2). The recorded force pulses show a non-monotonous fluctuation, indicating a stimulus-dependent instability. The instability in the *in-vivo* experiment, *ξe*, is defined as the standard deviation versus the average value of the force amplitude:

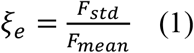

An example of *ξe* curves is shown in Figure 1(e3) by stimulating the nerve with a range of stimulus amplitudes. We identified a stimulus-dependent *ξe* pattern in terms of the *ξe* peak amplitude and position. This stimulus-dependent *ξe* is the major focus of this study.

### 3. Modeling instability by Circuit-Probability theory

The result of electrical stimulations can be modeled using the Circuit-Probability theory (C-P theory) ^[46]^. The major principle of the C-P theory can be explained by:

1. The electric field (E-field) across the axon membrane evokes action potentials. Since the phospholipid bilayer of a cell membrane can be modeled as a capacitor, this E-field is proportional to the cross-membrane potential, which is the voltage upon the capacitor of the cell membrane.
2. Since the input current and the generated voltage on the cell membrane do not share the same waveform, we need to build a circuit to calculate the voltage waveform.
3. There is a threshold voltage of nerve stimulation. Therefore, only the part of the voltage waveform exceeding the threshold has a probability of evoking the action potential.

A brief illustration of the C-P theory is proposed in Figure 2. It is known that a successful electrical stimulation of an axon will activate an action potential (Figure 2(a)). However, if the stimulus is insufficient to evoke an action potential, an oscillating voltage waveform, which is normally called subthreshold oscillation, will be recorded in the experiment of patch-clamp (Figure 2(a)). This subthreshold oscillation is the voltage applied to the cell membrane. Thus, the part of the voltage higher than the threshold shall have a probability of evoking an action potential. It is emphasized that the subthreshold oscillation may not be really subthreshold. There is still part of the voltage exceeding the threshold to provide a certain probability of activating an action potential. If the activation is failed, the recorded voltage is called subthreshold oscillation.

**Figure 2.**
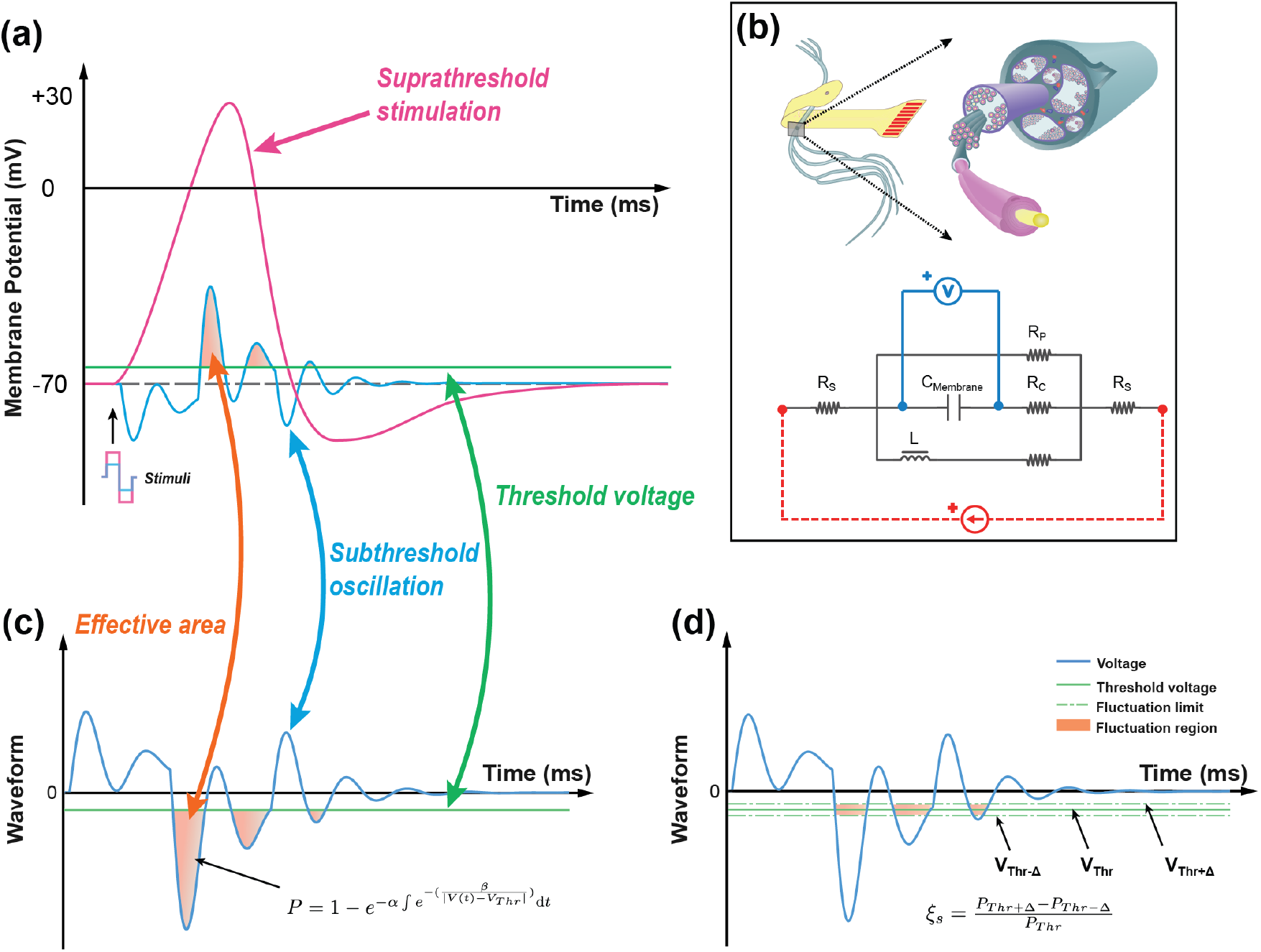
The concept of the C-P theory and the definition of instability in modeling. (a) The neural response of electrical stimulations: a suprathreshold stimulation will generate an action potential, while a subthreshold stimulation will induce a subthreshold oscillation. (b) The equivalent circuit with an RLC configuration to model the neurons. (c) The concept of C-P theory. The subthreshold oscillation can be duplicated by the voltage response of the RLC circuit in (b). The part of the voltage exceeding the threshold will be involved in the calculation of the probability for generating an action potential. (d) The definition of the instability of electrical stimulations: assuming that the threshold will have a fluctuation in the range of ±∆, the instability in modeling is defined as 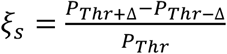.

To calculate this probability, we need two components. One is the equivalent circuit to duplicate the subthreshold oscillation. Another is the probability calculation based on the oscillating voltage. Our previous study demonstrated that this oscillating voltage could be duplicated by an RLC circuit shown in Figure 2(b) ^[46]^. In the circuit component, we build an RLC circuit to represent the passive property of neural tissue (Figure 2(b)). The capacitor *C*_*MeMbrane*_ represents the cell membrane. This circuit has an inductor, which differs from the RC circuit used in conventional models such as the Hodgkin-Huxley model ^[47]^. The origin of this inductor comes from the spiraling structures of myelin sheaths and the flexoelectricity effect of the cell membrane, which has been explained in our previous work ^[48]^.

Figure 2(c) shows a typical subthreshold oscillation generated by the RLC circuit in Figure 2(b). This voltage oscillation was reported in many biological experiments ^[49–61]^, particularly in the original study of the Hodgkin-Huxley model ^[47]^. When the threshold voltage is applied, the part of voltage exceeding the threshold will be involved in the equation of probability calculation:

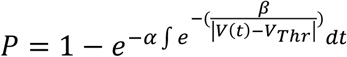

This equation is proposed and explained by our previous work on Circuit-Probability theory ^[46]^. α, β are parameters to be tuned to fit the experimental data. *V*_*Thr*_ is the threshold voltage, which is the difference between the resting potential and threshold voltage in Figure 2(a). Due to the voltage oscillation, more than one block of the voltage waveform can be involved in the probability calculus. In the case shown in Figure 2(c), three blocks are involved.

Since *V*_*Thr*_ is the difference between the resting potential and the threshold voltage in Figure 2(a), the fluctuation of the membrane potential, which is reported as the origin of the electrical nerve stimulation instability ^[9–11]^, can be modeled with a fluctuation of *V*_*Thr*_, as shown in Figure 2(d). We assume that the threshold voltage can only fluctuate within a region defined by Δ, then the threshold voltage can be changed from the upper limit *V*_*Thr*_ + ∆ to the lower limit *V*_*Thr*_− ∆, shown in Figure 2(d). The instability in the simulation can be defined as

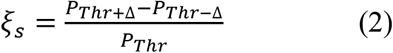

where ξ_*s*_ refers to the instability in simulation. *P*_Thr_, *P*_Thr+∆_ and *P*_Thr−∆_ are the calculated probability by setting their own threshold voltages.

A typical result of ξ_*s*_ by applying a positive-first biphasic square wave current with 400 µs pulse width is shown in Figure 3 (a). The curve has three peaks, reproducing the pattern shown in Figure 1(e3). Here we need to explain why our model can reproduce the pattern of peaks in *in-vivo* tests.

**Figure 3.**
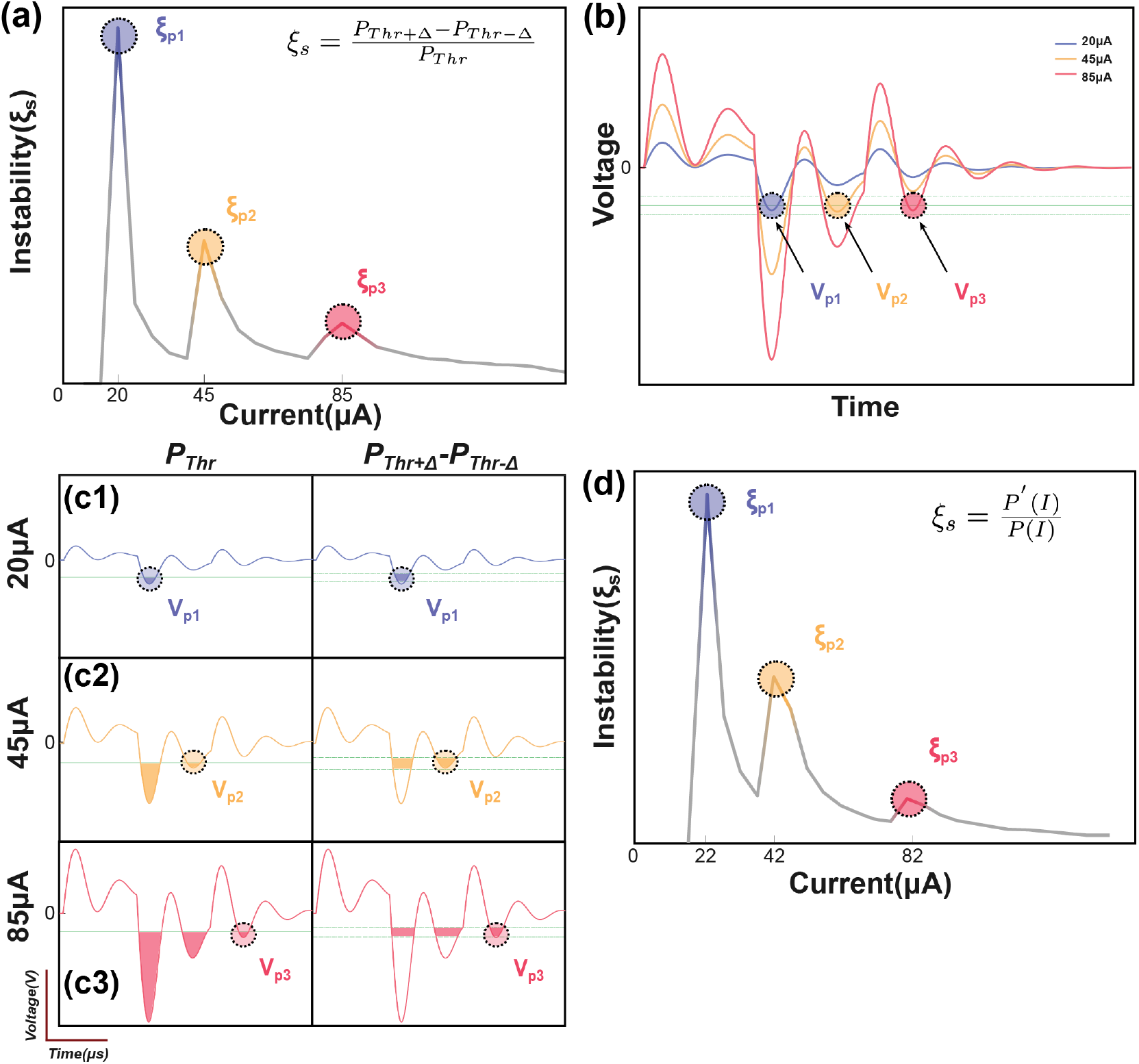
The illustrative explanation of the instability peaks in modeling. (a) Modeling results show several instability peaks in the instability curve by the definition of 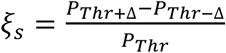 The voltage oscillation shows several oscillation peaks corresponding to the instability peaks in (a); (c) Illustrative explanation about how the voltage oscillation peaks induce instability peaks; (d) The modeling of instability peaks by the definition of 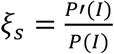.

For the representative voltage waveform shown in Figure 3 (b), three oscillation peaks can exceed the threshold, named as *V*_p1_, *V*_p2_ and *V*_p2_. Since these three voltage peaks are of different amplitudes, they reach the threshold in series by increasing the current.

When the current is low, only *V*_p1_ reaches the threshold (Figure 3(c1)), the instability is

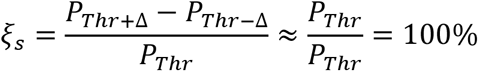

In this scenario, the instability reaches the maximum, shown as the first peak, ξ_p1_, in Figure 3(a). Then the instability will decrease by increasing the current until the second voltage peak, *V*_p2_, reaches the threshold, shown in Figure 3(c2). It will also induce a local maximum, ξ_p2_. But because the total probability, *P*_Thr_, on the denominator is higher, the amplitude of the second instability peak, ξ_p2_, is lower than ξ_p1_. Then by further increasing the current to make *V*_p3_ reach the threshold (Figure 3(c3)), a third instability peak, ξ_p3_, will appear, and its amplitude is lower than ξ_p2_. In summary, an instability peak will appear with a decreased amplitude whenever a new voltage peak exceeds the threshold.

The definition of instability, ξ_*s*_ in Figure 3(a) is based on the upper and lower limit of the threshold fluctuation, Δ. The amplitude of Δ will not change the qualitative results, which refer to the number and position of those peaks of ξ, but will determine the quantitative result, which refers to the height of the peaks. Here we will conduct a further derivation to make the definition of ξ free of Δ.

The fluctuation of the threshold changes the area of the voltage waveform involved in the probability calculus. Meanwhile, changing the amplitude of the input current can induce the equivalent effect. For example, increasing the threshold voltage is equivalent to reducing the current amplitude, while decreasing the threshold voltage is equivalent to increasing the current amplitude. Thus, equation (2) can be rewritten into another form:

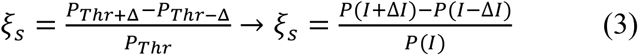

In this new equation (3), the fluctuation of the threshold is replaced by a change in the input current.

Since

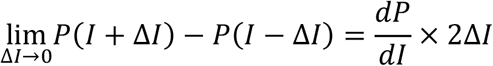

Then further derive the equation (3) as follow:

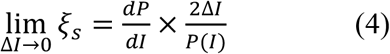

Since ∆*I* is always set as a constant, it can be neglected in the qualitative modeling; the equation (4) can be written as follow:

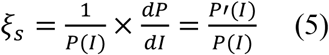

This new definition does not rely on the value of Δ. The instability curve in Figure 3(d) modelling by equation (5) can generally reproduce the pattern in Figure 3(a) modelling by equation (2). But the width and the position of the peaks will have a slight shift, which is induced by reducing Δ to zero. In the following sections, all simulation results used equation (5) as the definition of ξ_*s*_.

## Results

### 1. The number and position of peaks

Based on the stimulus-response relationship given by the C-P model, we generated a full mapping of the instability by traversing all possible values of the current amplitude and the pulse width while keeping the same waveform. The simulation results will form a heat mapping shown in Figure 4(a), called an instability mapping. All modeling settings and parameters are listed in Table 2. The result of the instability curve of a specific pulse width is just the profile of one cross-section of the instability mapping. In Figure 4(a), the profiles captured at different pulse widths show different patterns (Figure 4(b)). This changing trend can also be observed in *in-vivo* testing (Figure 4(c)). Our model closely reproduces the observed patterns *in vivo*.

**Figure 4.**
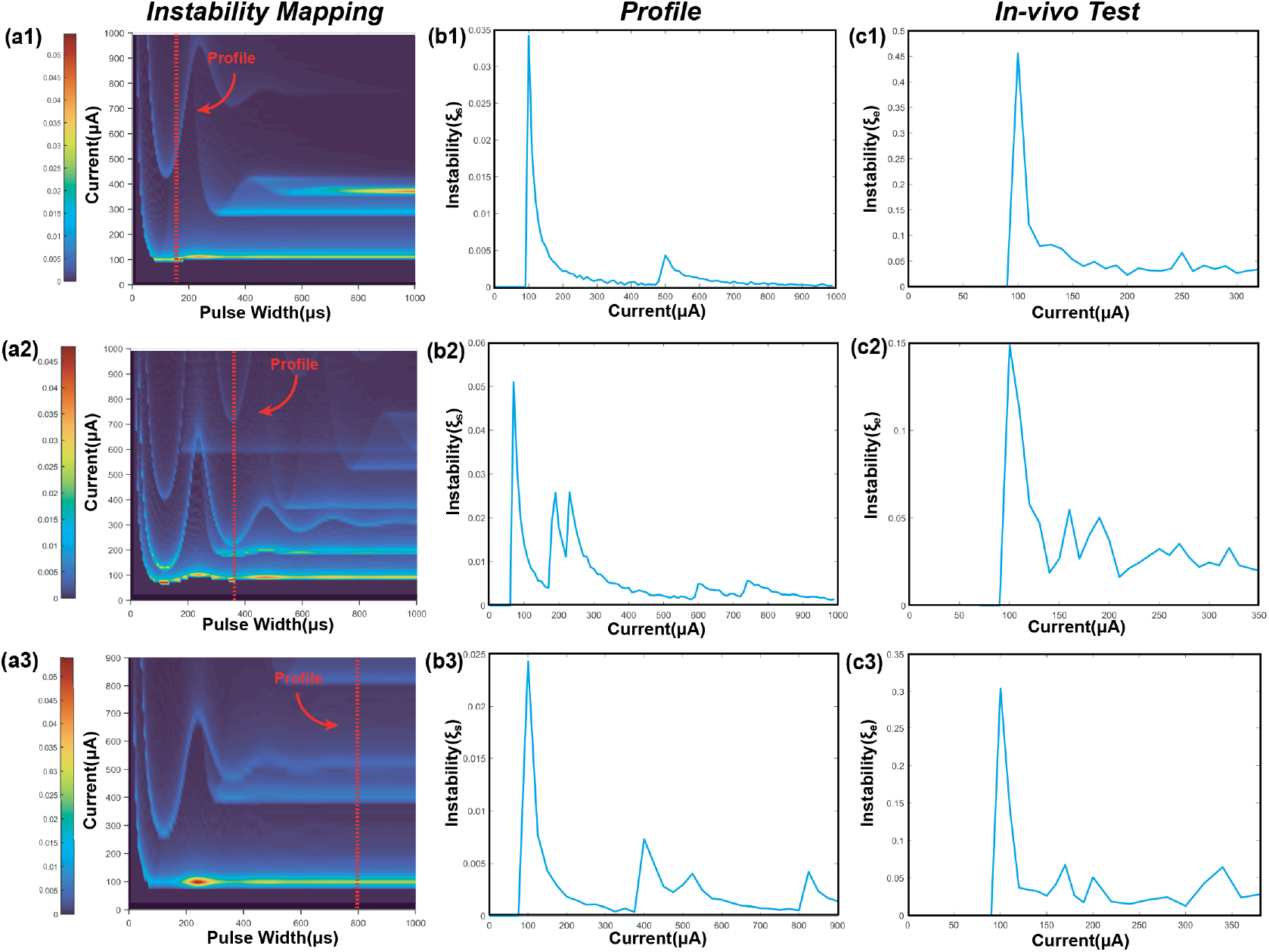
The patterns of the instability peaks in *in-vivo* tests are reproduced by modeling. (a1-a3) The heat map of the instability by changing both pulse width and current amplitude; (b1-b3) The captured profile of the instability curve at a specific pulse width; (c1-c3) The measured instability curves to be reproduced by (b1-b3).

### 2. The moving track of the peaks

It is clearly observed that each peak moves along a specific track in the instability mapping. This moving track can also be reproduced by our modeling results shown in Figure 5. In Figure 5(a1), we tune the modeling parameters to generate an instability mapping to fit the *in-vivo* results shown in Figure 5(b1).

**Figure 5.**
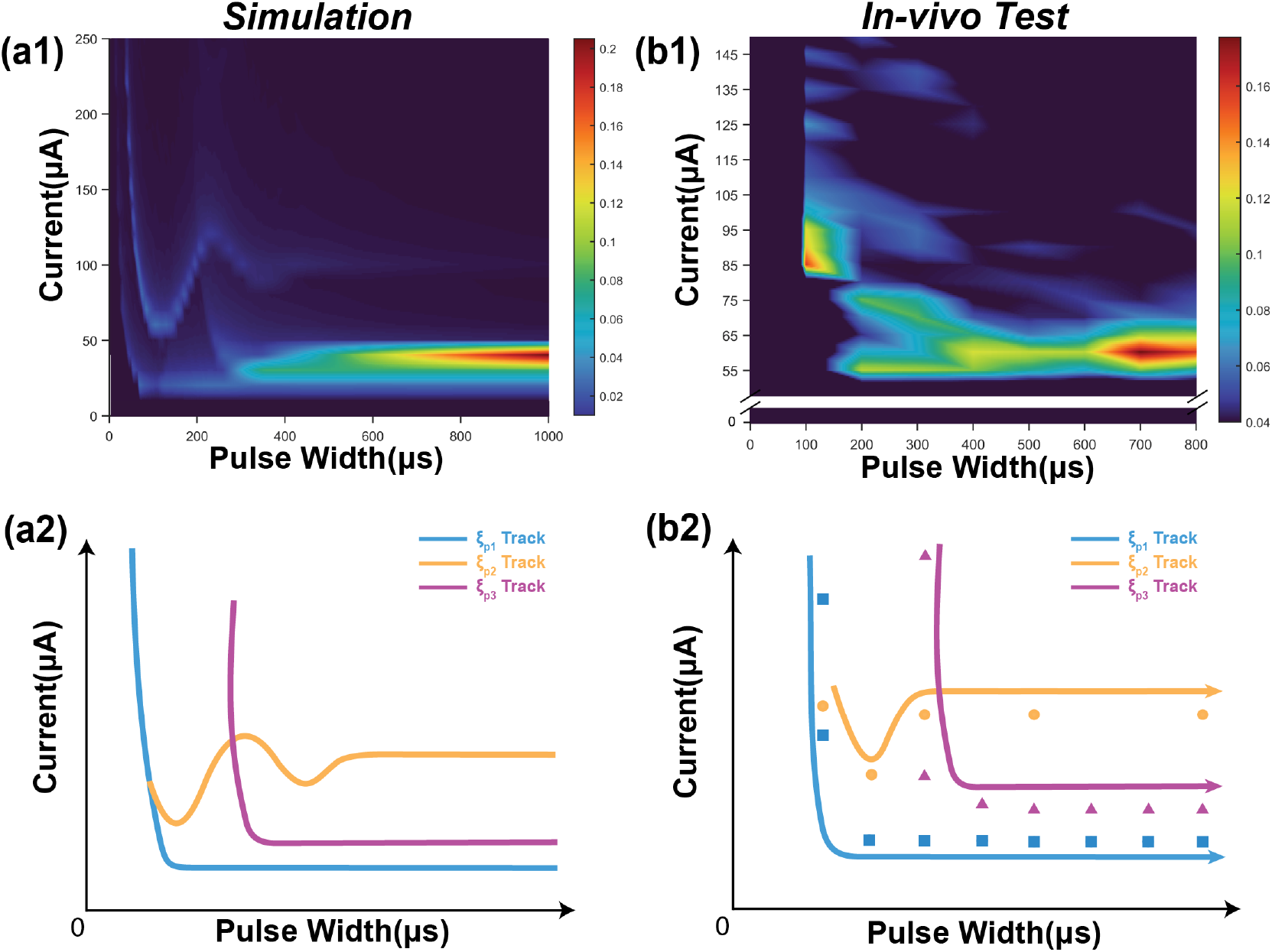
The moving tacks of instability peaks in *in-vivo* test are reproduced by modeling. (a1) The heat map of the instability by modeling; (a2) The tracks of the three instability peaks in (a1); (b1) The measured heat map of the instability by *in-vivo* test; (b2) The recognized positions and tracks of the instability peaks in (b1).

In Figure 5(a1), three peaks can be found, named as ξ_*P*1_, ξ_*P*2_ and ξ_*P*3_. Their moving tracks are labeled in Figure 5(a2). The moving tracks of ξ_*P*1_ and ξ_*P*2_ share a similar “L” shape. They have a large divergence at the low pulse width and tend to merge at the high pulse width. The moving track of ξ_*P*3_ has an individual shape with some oscillation.

The in vivo results in Figure 5(b1) can generally be fitted by the modeling results in Figure 5(a1). The data to generate the heat map in Figure 5(b1) is a sparse matrix, showing key features of the instability mapping. We labeled the position of the peaks of *in-vivo* testing in Figure 5(b2). Based on the guidance of Figure 5(a2), it is found that the positions of the peaks also closely fit the predicted moving tracks. The modeling parameters can be found in Table 2.

Since the modeling parameters mainly determine the instability mapping generated by C-P theory, we also generate some other possible patterns of the instability mapping by changing the circuit parameters. In this study, we only show how the patterns change with the quality factor, 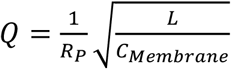, and the resistor connected in series with the inductor, RL.

In Figure 6, the quality factor, Q, mainly determines the spacing between the moving tracks of the peaks. A low *Q* factor will increase the spacing, while a high Q factor will make the tracks closer to each other. The resistor RL mainly determines the density of the peaks. A lower RL generates more peak tracks.

**Figure 6.**
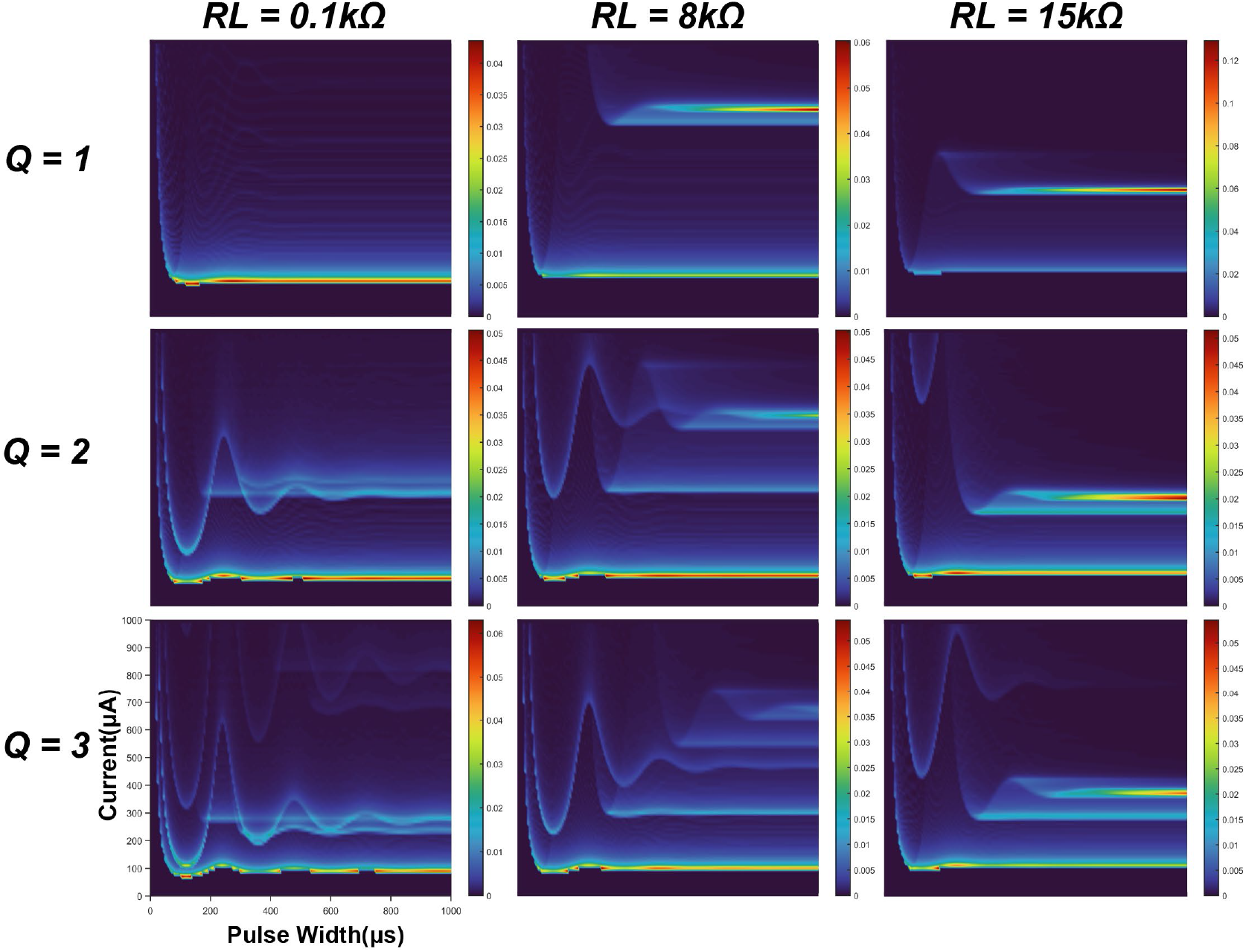
A parameter comparison by modeling. The resistor connected in series with the inductor and the quality factor of the circuit are selected as the variables.

The patterns have quite complex changing trends, which cannot be fully elucidated here. Figure 6 only demonstrates a facet of the effect of the parameters on the modeling results, showing that the patterns and tracks of the peaks are highly tunable. The modeling parameters can be found in Table 2.

### 3. How to improve the stability of electrical nerve stimulations by parameter selection

Our modeling work also provides a possible method to optimize the stimuli parameters to improve the stability of electrical stimulations. We make a case demonstration based on the modeling results in Figure 5(a1), which is replotted in Figure 7(d).

**Figure 7.**
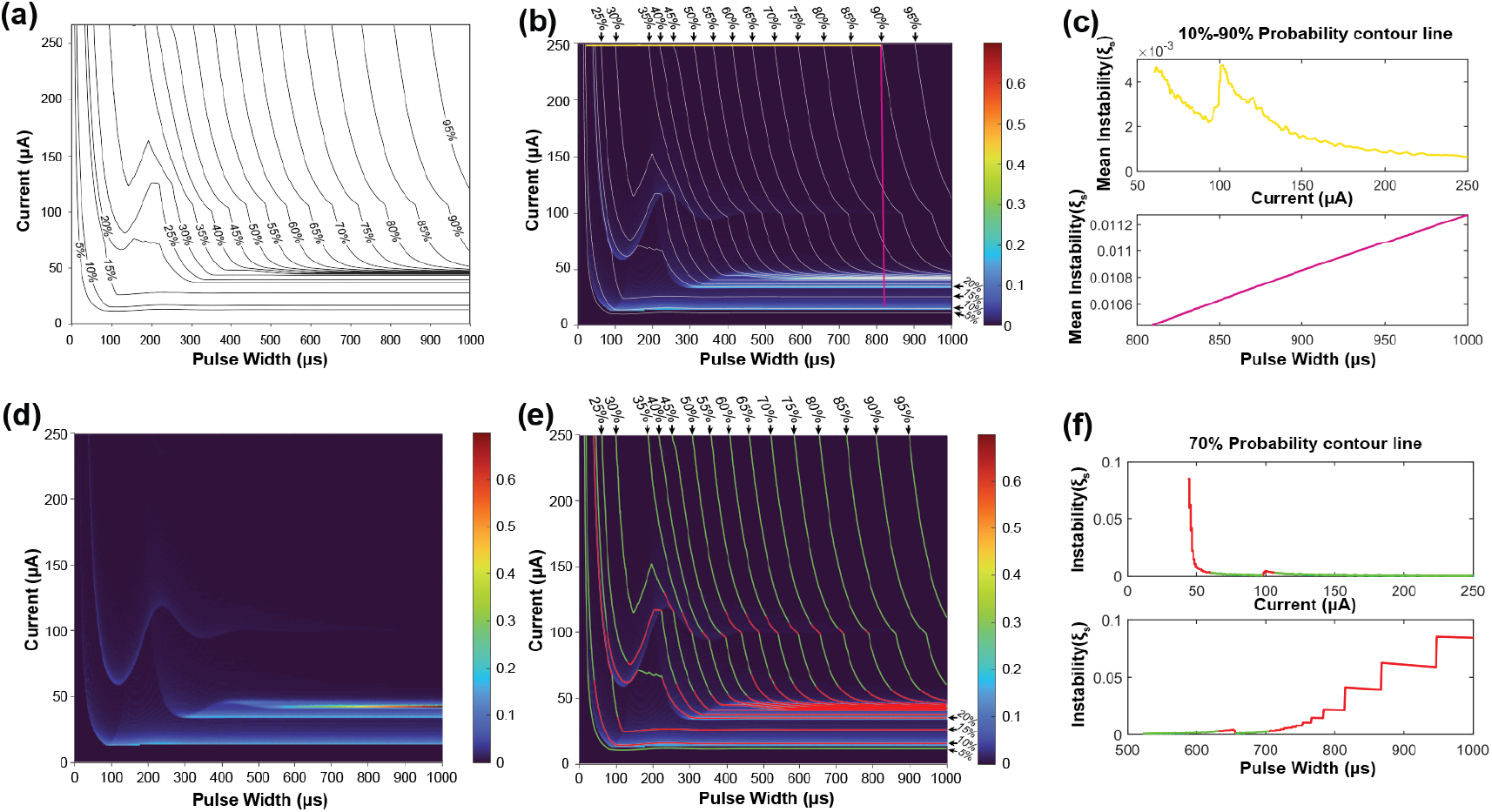
The methods for improve the stability of electrical nerve stimulations by selecting proper stimuli parameters. (a) The contour map of the probability based on the modeling results in 5(a1). (b) An overlapping of the contour map and its instability mapping. The horizontal yellow line is the parameter path with the minimal total instability in all horizontal lines; the vetical purple line is the parameter path with the minimum total instability in all vertical lines. (c) The calculated total instability of all horizontal lines (yellow) and vertical lines (purple). (d) A duplicated instability mapping as in Figure 5(a1). (e) An overlapping of the contour map and its instability mapping with green region (low instability) and red region (high instability). (f) A case demonstration of the instability calculation of the 70% contour line by setting current (upper) and pulse width (lower) as x-axis. The curve higher than ξ_*Threshold*_ is labeled in red, while the curve lower than ξ_*Threshold*_is labeled in green.

Since the C-P theory can generate the whole probability mapping for all stimuli parameters, we can also generate a contour map of the probability, as shown in Figure 7(a). Improving stimuli parameters is to find a path from a very weak stimulation (10%) to a very strong stimulation (90%) on the map (Figure 7(b)) with the lowest total instability.

Firstly, we demonstrate the improving method by a simple parameter selection, which is fixing either the current amplitude or the pulse width. In this case, the path of the parameter in Figure 7(b) should be either a vertical straight line (fix the pulse width) or a horizontal straight line (fix the current amplitude). All the data points of the instability on each line are summed up for comparison, and the results are shown in Figure 7(c). For the situation of fixing the current, the calculation results are shown as the yellow line in Figure 7(c). This data curve shows some fluctuation and reaches the minimum at the current of 250 µA with total instability of about 0.001, which refers to the yellow horizontal line at the top of Figure 7(b). For the situation of fixing the pulse width, the results are shown as the purple line in Figure 7(c). It is a monotonous increasing line with the pulse width. Thus, the minimal instability happens at about 810 µs with total instability of about 0.0104, which refers to the purple line in Figure 7(b). As seen, the instability of the purple line is about 10 times higher than that of the yellow line. The reason is that the vertical purple line inevitably goes through the region with very high instability (indicated with red and yellow colors, named as high instability region) at the bottom of Figure 7(d), while the yellow line is located at the top of Figure 7(b), which is a region with low instability. As seen, the key to improving the stability is to avoid the high instability region. Meanwhile, the modeling results in Figure 6 show that the high instability regions normally distribute horizontally. Thus, horizontal lines have a better chance of completely avoiding these regions. On the contrary, if the horizontal line accidentally across one of the high instability regions (which may happen in applications), all stimulations will exhibit high instability. Thus, it is recommended that a proper instability mapping, as in Figure 5(a1), should be characterized to indicate the high instability regions. Just by avoiding these regions, the stability of electrical nerve stimulations can be significantly improved.

Apparently, if the parameter path is not a straight line, or even not a continuous path, it is possible to avoid all high instability regions. To demonstrate this method, the instability of along each contour line is calculated. As a case demonstration, the calculation results of the contour line of 70% are shown in Figure 7(f) by setting pulse width and current as x-axis, respectively. As seen, even with the same probability, the instability can have a dramatical difference by changing the stimuli parameters. Here we give a rule for setting the threshold for stimuli parameter selection:

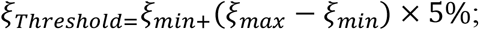

Here ξ_*max*_ and ξ_*min*_ refers to the maximum and minimum instability on the map, respectively. The region higher than ξ_*Threshold*_ is indicated with red color, while the region lower than ξ_*Threshold*_ is indicated with green color. As seen in Figure 7(e), the green regions on the contour lines can roughly form the area, which parameter path can go though with very low instability.

This method may be quite difficult to be applied in applications, since a fine characterized instability mapping is very difficult and time consuming. The key inspiration of this modeling is that, even with the same probability, which refers to the stimulation strength, the instability can vary a lot by setting different parameters. A proper parameter selection can significantly improve the stability of electrical nerve stimulations.

## Discussion

### 1. The effect of the inductor involved in the equivalent circuit of neural tissue

The most controversial part of the C-P theory is the inductive factor involved in the equivalent neural circuit. Starting from the cable theory and H-H model, a neural circuit of RC configuration is always applied to model the axon. However, the passive voltage response does follow an RLC circuit, which has been validated by our previous works ^[46,48]^ and many previous studies ^[49–61]^. To further confirm the effect of the inductor on the instability in electrical nerve stimulation, the modeling results using an RC circuit (Figure 8(a)) are shown to make a comparison. A typical RC voltage response by applying a square current pulse is shown in Figure 8(b). Due to the lack of oscillation, only one voltage block can exceed the threshold. Thus, the calculated probability shows a smooth increasing curve without any abrupt change (Figure 8(c)). Therefore, only one instability peak will be observed in modeling results (Figure 8(d1-d3)). The track of the instability peak will form an “L” shape, which is not affected by modeling parameters. So based on our theory, if the neural circuit really follows an RC configuration, there shall always be only one peak, which is inconsistent with the experimental data. Meanwhile, compared with the simple peak track in Figure 8(d), the instability mapping in Figure 5(b1) shows a much more complex pattern of the tracks. Therefore, an RLC circuit can reproduce the passive property more accurately than an RC circuit.

**Figure 8.**
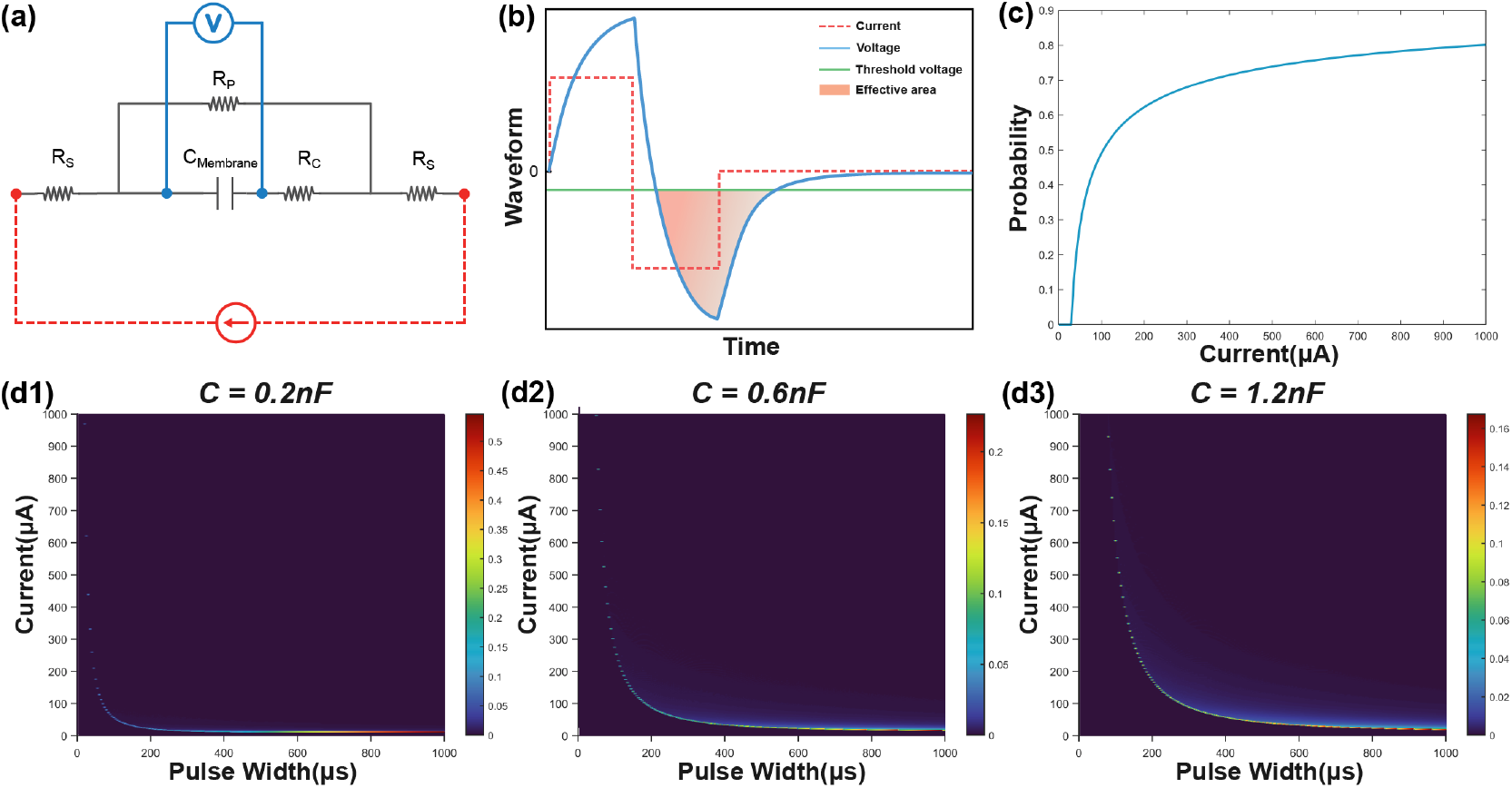
A comparison of the modeling by using an RC circuit. (a) The RC circuit used in modeling; (b) A representative voltage waveform (blue line) by applying a square current (red dash line). Only the voltage (filled with orange color) exceeding the threshold (green line) is calculated in probability calculus. (c) The probability is calculated by changing the current amplitude; (d1-d3) The heat map of the instability by changing the value of the capacitance.

### 2. The effect of current waveforms

Our model suggested that the instability is also affected by the shape of the current waveform. To demonstrate this, two current waveforms, square wave and sine wave, were involved in the test shown in Figure 9. Figure 9(a) shows the measured force by changing the current amplitude. The applied pulse width is 200 µs. The calculated instability is shown in Figure 9(b). We set the force as the x-axis, which provides a fair basis for the comparison (Figure 9(c)). Our results indicated that the sine wave induced a higher instability. It is worth investigating more with a wide range of stimulation waveforms and parameters in future studies.

**Figure 9.**
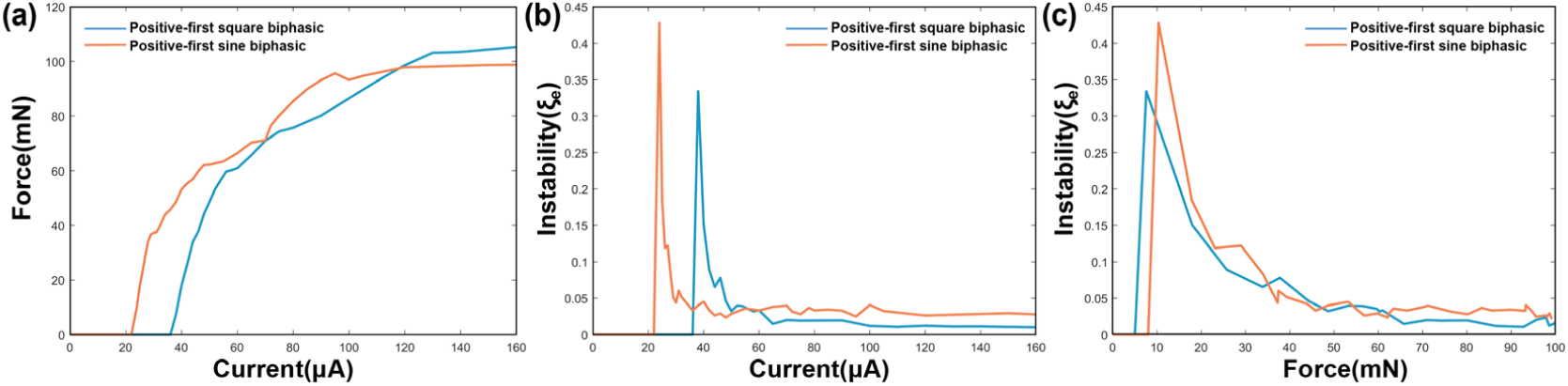
A comparison between square and sine current waveform. (a) The measured force; (b) The calculated instability curve; (c) Re-processed instability curves by setting the force as x-axis.

### 3. Instability versus linearity

It is expected to minimize the instability of the electrical stimulation. Meanwhile, it is also expected to achieve a linear control of the stimulation strength. However, based on our theory, there is a trade-off between these two factors. Based on the definition of instability,

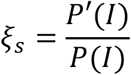

It can be derived that

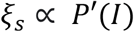

While

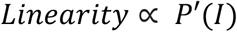

Thus,

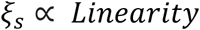

It means that the instability tends to be high in the range of electrical stimulation, which can provide a good linear relationship between the current amplitude and the neural response. While in the range that the strength of the neural response is not much affected by the current amplitude, the stimulation will be more stable. Thus, there is a trade-off between stability and stimulus settings.

A more illustrative explanation of this trade-off is shown in Figure 10. Figure 10(a1) shows the probability curve by changing the current amplitude, which can generally duplicate the testing results in Figure 10(b1), the measured force by changing the current amplitude. The model parameters can be found in Table 2. In Figure 10(a1), the probability curve is not smooth. Besides the beginning section, there are another two positions with abrupt changes labeled as circles with different colors. The derivatives of the probability curve at the circles reach the local maximum, corresponding to the three peaks in the instability curve (Figure 10(a2)). In Figure 10(b1), similar abrupt change points can also be found. The instability curve in Figure 10(b2) suggests that the peaks happen at the position of these abrupt changes. More similar data can be found in Supplementary Figure S2.

**Figure 10.**
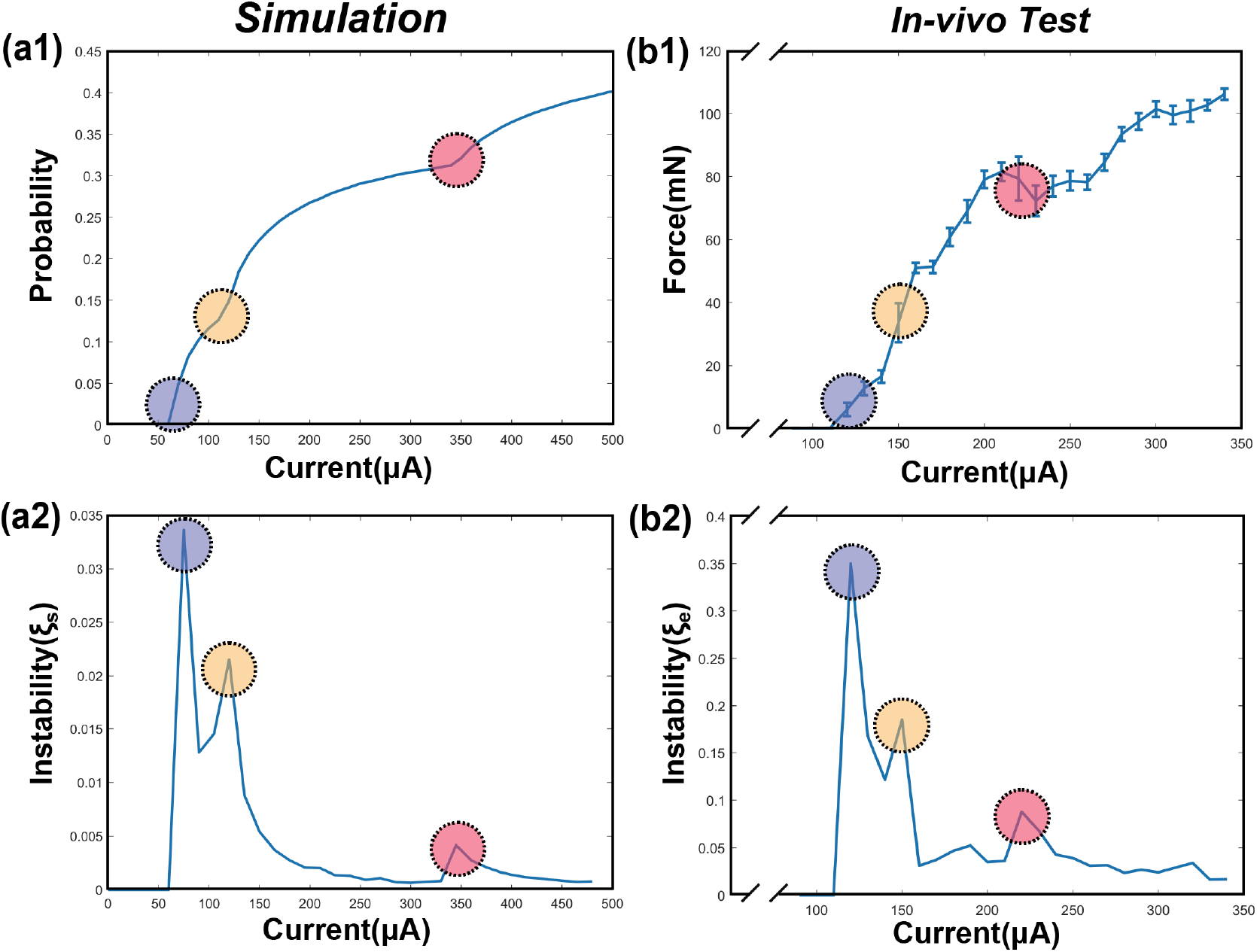
The modeling and in-vivo results show the trade-off between electrical stimulations’ stability and linearity. (a1) A representative probability curve of modeling showing several abrupt change points; (a2) The calculated instability curve shows the positions of the peaks corresponding to the positions of the abrupt change points; (b1) A presentative force curve of the *in-vivo* test showing several abrupt change points; (b2) The instability curve shows the positions of the peaks corresponding to the positions of the abrupt change points.

This study first proposed and validated the trade-off between stability and linearity in electrical stimulation. The prediction and explanation of this trade-off show that our theory does not only fit the testing data but also elucidates its mechanism.

### 4. The issue of historical path divergence

We proposed a new concept called historical path divergence. It is essential to understand the phenomenon of continuous electrical nerve stimulation. A brief illustration of this concept is shown in Figure 11.

**Figure 11.**
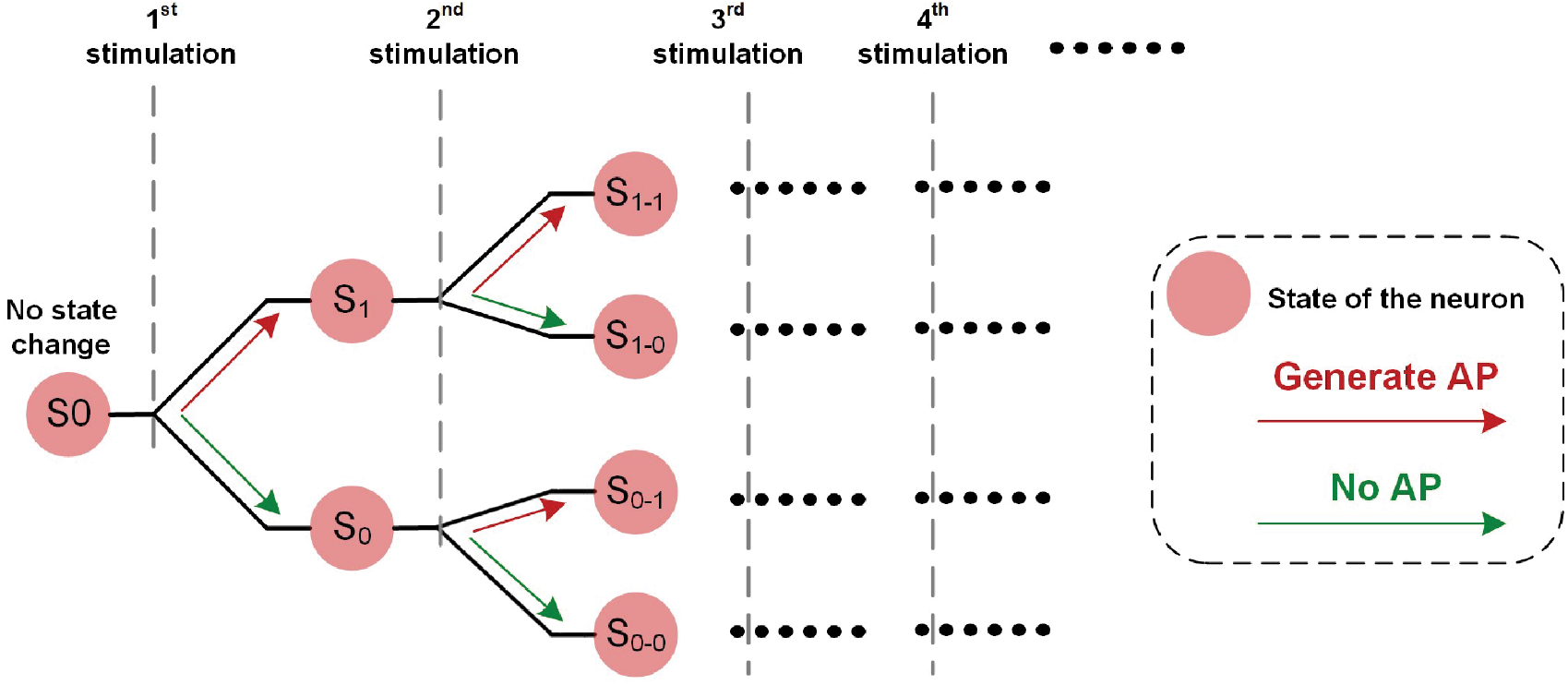
The illustrative explanation of the historical path divergence issue.

In Figure 1(e3), the fluctuation of the measured force demonstrated a non-monotonic trend. So, it can be inferred that the state change of the neuron, which refers to the threshold fluctuation, is determined by both the electrical input and the real-time neuron states.

At the beginning of the electrical stimulation, all neurons are at state S0, which means no state change. Since all neurons are in the same state, they are synchronized. After the first stimulation is applied, two possible outcomes might occur: S1 (generate AP) and S0 (no AP). Since the number of possible states for all neurons increases, they are less synchronized or more asynchronous (that is, neurons tend to have different states). Each state can have two possible outcomes when the subsequent stimulation is applied. Therefore, the number of possible states is increased, causing decreased synchronicity of neurons.

The stimulus-induced historical paths formed a binary tree in Figure 11. The synchronicity of the states of all neurons will determine the observed instability. When the synchronicity is higher, such as state S0, all neurons tend to have the same state change post-stimulus. This state change can either increase or decrease excitability. Therefore, the observed force will have a more evident fluctuation trend, either increasing or decreasing. With subsequent stimulations, neurons trends to be asynchronous. Some may increase, and some may decrease. Thus, the observed force fluctuation will not have an evident changing trend. In other words, the measured force can be more stable.

At the beginning of the stimulation, the synchronicity across neurons is high. Thus, the resulting force tends to have a more evident changing trend with more increased instability. Meanwhile, this evident changing trend is the same in all testing trials. However, with subsequent stimuli, this evident changing pattern tends to disappear, and the force tends to be more stable.

The abovementioned conclusions are validated by the *in-vivo* experiments shown in Figure 12. Figure 12(a) shows the curves of five testing trials with the same stimulation parameters. The standard deviation (SD) can represent the instability of the curve. To show the change in the SD with time, the SD is calculated with each 5 data points (Figure 12(b)). SD is high at the beginning and soon decreases to a certain level, showing the trend that the force becomes more stable with time. Figure 12(c) only shows the first 15 data points of the force curve in Figure 12(a). The changing trend is labeled with colors (green refers to decrease, and red refers to increase). Only at the beginning stage (the first 4 data points), all force curves show the same decreasing trend. The data analysis in Figure 12 shows the effect of the historical path divergence. Continuous stimulation will diverge the historical paths of all neurons and reduce instability. Thus, instability is always the highest at the beginning stage.

**Figure 12.**
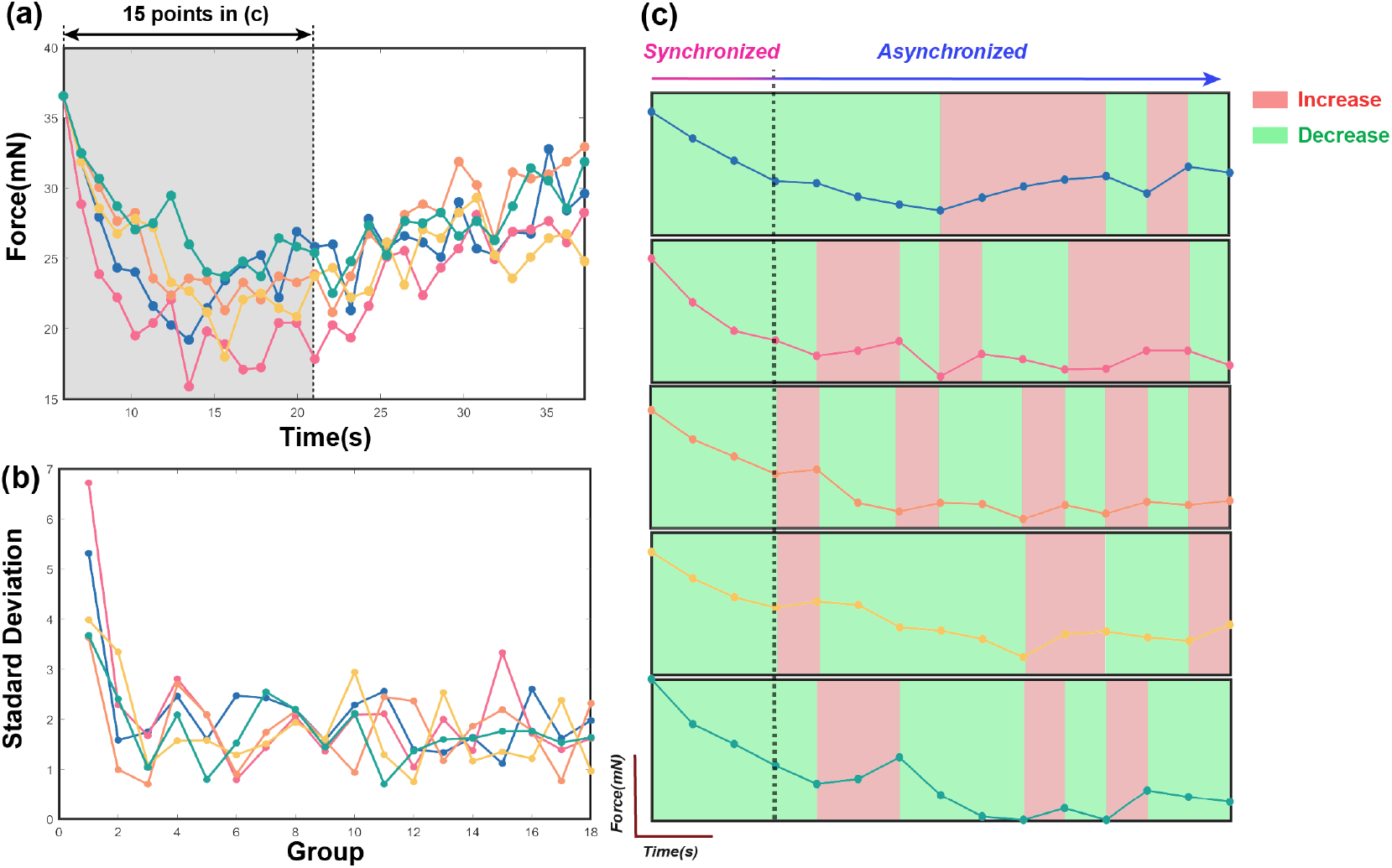
The testing data to validate the historical path divergence issue. (a) The five measured force curves with the same testing parameters; (b) The standard deviation of the force curve in (a). Every five points in (a) are calculated as one data point in (b). (c) The first fifteen points of the five curves in (a) are compared. The changing trends are labeled in colors (red refers to increasing, and green refers to decreasing).

## Limitations

Although we can reproduce the patterns of peaks in the modeling results, the heights of the peaks are not accurate. Several factors might limit the modeling accuracy.

### 1. The parameters in the C-P theory are not accurate

We cannot precisely assign the parameters in C-P theory. Our previous work developed the C-P theory to build the mathematical relation between the electrical input and the neural response, which is similar to the H-H model. However, unlike the H-H model, the C-P theory can describe macroscopic electrical nerve stimulation. Thus, those parameters involved in the model are not measurable. Meanwhile, due to the high nonlinearity of the model, it is also difficult to precisely estimate these parameters. Currently, we assign the value of each parameter by the exhaustive method, which can roughly reproduce the distinctive features of the testing results, such as the number and position of the peaks.

### 2. The value of threshold fluctuation is unknown

The instability’s origin is known as the membrane potential fluctuation. However, it is difficult, if not impossible, to measure the value and range of the membrane potential fluctuation in vivo. This value will quantitatively affect the height of the peaks in our modeling results but does not affect the general patterns. Therefore, we give a new definition of instability free of the value of membrane potential fluctuation *P*′/*P*’, which is more suitable for qualitatively reproducing the patterns of the peaks.

### 3. The definitions of instability of *in vivo* testing and modeling are not consistent

The instability of in vivo testing is based on the measured force’s standard deviation (SD). However, the SD cannot be generated in our modeling. Instead, we used the difference between the maximum and minimum probability, which positively correlated with the SD. This inconsistency will not affect the reproduction of the patterns but will forbid us from acquiring the accurate height of peaks.

## Conclusion

There is much-growing evidence that the performance of electrical stimulation cannot be significantly improved by merely optimizing stimulus parameters without considering the complexity of the biophysical characteristics of the target nerve system. We investigated the characteristics of the instability of electrical nerve stimulation by the Circuit-Probability theory. Our model reveals several critical characteristics of the instability peaks. Firstly, the instability peaks’ physical origin is the axon’s oscillatory nature, whose passive electric property shall be modeled by an RLC circuit. Thus, due to the complexity of the RLC circuit’s voltage response, the measured instability peaks will follow certain patterns which our model can reproduce. On this basis, it is possible to improve the stability of electrical nerve stimulation by a proper stimuli parameter selection. A case demonstration with different parameter selection methods is provided. Meanwhile, the measured pattern of the instability peaks substantially supports the inductive factor in neural circuits. Moreover, our model reveals the trade-off between the stability and linearity of electrical nerve stimulations. That is, given a current range that allows linear control of stimulation strength, the stability of neural response tends to be low. Finally, our model proposes a new perspective to study electrical nerve stimulation: the historical path divergence that predicts the instability of the electrical nerve stimulation changes with the stimulation duration.

## Acknowledgment

This work was supported by the grants from Guangdong Research Program (2019A1515110843), Shenzhen Research Program (GJHZ20200731095206018, JCYJ20210324101610028, JCYJ20180507182057026) and National Natural Science Foundation of China grants (31800871, 82171082, 62071459) and National Key Research and Development Program of China (2022YFF1202500, 2022YFF1202502). Hao Wang proposed the theory. Shoujun Yu carried out the modeling process. Shoujun Yu, Yonghong Liu, Wenji Yue and Yapeng Zhang carried out the in-vivo tests. Sara Khademi and Wenji Yue fabricated the neural probes. Tian Zhou, Zhen Xu and Fenglin Liu helped train the experimental process. Tianruo Guo helped refine the theory and improve the writing. Bing Song, Tianzhun Wu and Xuefei Yu provided the general guidance of the study. All authors contributed to the reference collection and idea discussion.

